# Acoustically Targeted Chemogenetics for Noninvasive Control of Neural Circuits

**DOI:** 10.1101/241406

**Authors:** Jerzy O. Szablowski, Brian Lue, Audrey Lee-Gosselin, Dina Malounda, Mikhail G. Shapiro

**Author notes:** Correspondence should be addressed to M.G.S.

## Abstract

Neurological and psychiatric diseases often involve the dysfunction of specific neural circuits in particular regions of the brain. Existing treatments, including drugs and implantable brain stimulators, aim to modulate the activity of these circuits, but are typically not cell type-specific, lack spatial targeting or require invasive procedures. Here, we introduce an approach to modulating neural circuits noninvasively with spatial, cell-type and temporal specificity. This approach, called acoustically targeted chemogenetics, or ATAC, uses transient ultrasonic opening of the blood brain barrier to transduce neurons at specific locations in the brain with virally-encoded engineered G-protein-coupled receptors, which subsequently respond to systemically administered bio-inert compounds to activate or inhibit the activity of these neurons. We demonstrate this concept in mice by using ATAC to noninvasively modify and subsequently activate or inhibit excitatory neurons within the hippocampus, showing that this enables pharmacological control of memory formation. This technology allows a brief, noninvasive procedure to make one or more specific brain regions capable of being selectively modulated using orally bioavailable compounds, thereby overcoming some of the key limitations of conventional brain therapies.

## INTRODUCTION

Neurological and psychiatric diseases together affect over 35% of the adult population^1–3^, and often involve the dysfunction of neural circuits defined by specific spatial locations and cell types^4–8^. However, conventional pharmacological treatments for such diseases act throughout the brain, leading to significant side-effects. While invasive surgery is able to target specific parts of the brain for excision or electrical stimulation, it carries significant risks. Emerging therapies based on gene or cellular therapy are also typically delivered using surgical injections, often with limited spatial coverage and acting in an always-on fashion lacking temporal dose control. Here, we introduce an alternative approach to neuromodulation that delivers spatial, cell-type and temporal control without surgery. This approach – which we call acoustically targeted chemogenetics, or ATAC – achieves this performance by combining three recently developed technologies: focused-ultrasound blood-brain barrier opening (FUS-BBBO) for spatial targeting, adeno-associated viral vectors (AAVs) for delivery of genes to specific cell types, and engineered chemogenetic receptors for modulation of targeted neurons using orally bioavailable compounds (Fig. 1, a).

**Figure 1.**
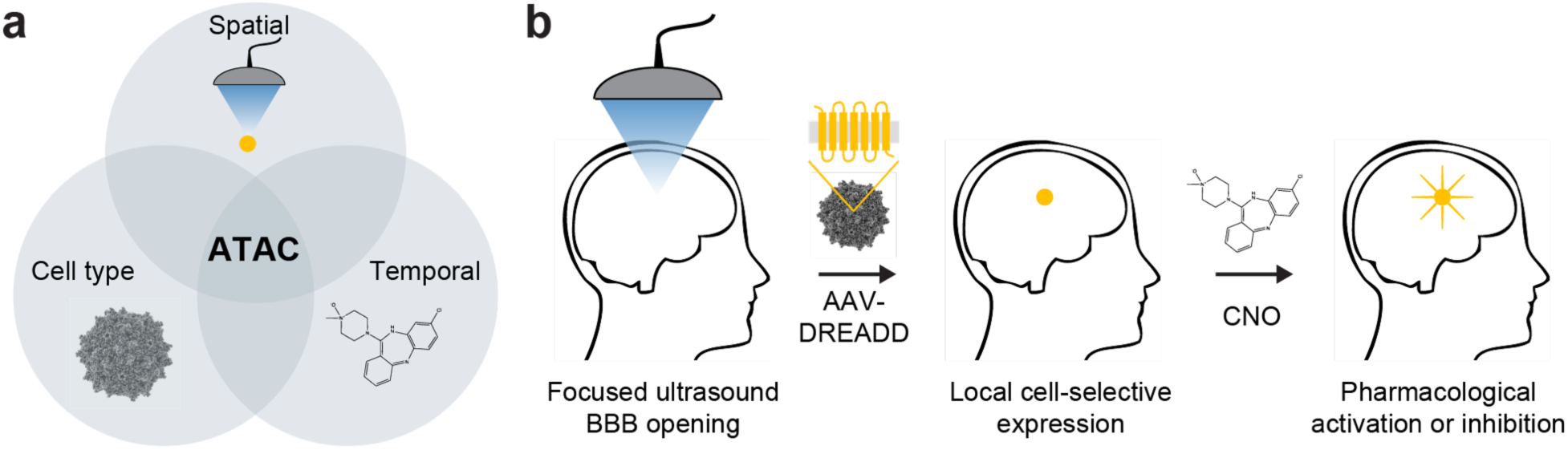
Acoustically targeted chemogenetics (ATAC) paradigm. (**a**) The ATAC paradigm provides a combination of millimeter-precision spatial targeting using focused ultrasound, cellular specificity using viral vectors with cell type-specific promoters driving the expression of chemogenetic receptors, and temporal control via the administration of the chemogenetic ligand. (**b**) In the ATAC sequence, the blood-brain barrier (BBB) is opened locally using focused ultrasound, and systemically injected adeno-associated virus (AAV) encoding a designer receptor exclusively activated by designer drug (DREADD) enters the treated area. After several weeks, the DREADD is expressed in the targeted region in cells possessing selected promoter activity. At any desired subsequent time, the DREADD-expressing neurons can be excited or inhibited through a chemogenetic drug such as clozapine-n-oxide (CNO).

FUS is an established biomedical technology that takes advantage of ultrasound’s ability to focus in deep tissues such as the brain with millimeter spatial precision^9–11^. FUS-BBBO combines transcranial ultrasound in the low-intensity regime with systemically administered microbubbles, whose stable cavitation in blood vessels at the ultrasound focus results in localized, temporary and reversible opening of the BBB^12,13^. This allows small molecules, proteins, nanoparticles or viral capsids^12,14–17^ to enter the brain at the site of applied ultrasound. FUS-BBBO has been used in multiple animal species^12,18,19^, and is currently undergoing clinical testing^9^. ATAC combines FUS with replication-incompetent AAV vectors, an established method to stably transfect mammalian cells without integration into the target genome. AAV is widely used in neuroscience research, and has recently shown promise in the clinic^20–24^. When a gene of interest carried by AAV is encoded under an appropriate promoter, the expression of this gene can be restricted to a specific class of neurons^25^. In ATAC, the AAV vector encodes chemogenetic receptors, a class of engineered proteins whose expression in neurons allows these cells to be controlled by otherwise inert systemically administered compounds^26, 27^. In particular, we use the Designer Receptors Activated Exclusively by Designer Drugs, or DREADDs. These proteins are modified versions of natural activatory or inhibitory GPCRs, engineered to respond to otherwise inert molecules such as clozapine-n-oxide (CNO) rather than endogenous ligands^28^.

In the ATAC paradigm, a one-time FUS-BBBO procedure “paints” the region or regions of the brain to be modulated, while AAV vectors and DREADDs sensitize specific neurons in these regions to subsequent excitation or inhibition with CNO (Fig. 1, b). While the three components underlying this paradigm have been separately established, and previous work has combined AAVs with either BBBO or DREADDs, the integration of these three technologies to achieve ATAC’s unique combination of spatial, cell-type and temporal control of neural circuits has not been reported. Here, we demonstrate the basic capabilities of ATAC in mice by evaluating the ability of this technique to selectively activate or inhibit excitatory neurons in the hippocampus and midbrain, regions involved in memory formation and volitional behavior and implicated in several neuropathologies. Our biochemical and behavioral experiments show that ATAC enables selective neuromodulation of these parts of the brain, and that inhibitory ATAC is able to reduce traumatic memory formation in a model of contextual fear learning.

## RESULTS

### Anatomical and genetic targeting of DREADDs

To evaluate the ability of ATAC to target the expression of DREADDs to a specific location in the brain, we first performed FUS-BBBO on the hippocampus of wild-type mice. The hippocampus is a brain region involved in memory formation and implicated in several neurological and psychiatric diseases, including anxiety, epilepsy and Alzheimer’s^29^. To achieve expression throughout this brain structure, we performed FUS-BBBO at 6 locations covering the ventral and dorsal hippocampus using an MRI-guided focused ultrasound instrument operating at 1.5 MHz (Fig. 2, a). FUS was applied immediately after an intravenous injection of microbubbles and viral vector, with a gadolinium contrast agent co-administered to visualize regions with successful BBBO. Each of the 6 targeted regions had an ovoid shape with a diameter of approximately 1 mm and a length of 4 mm (Fig. 2, b). As our viral vector, we chose AAV9, a serotype of AAV with favorable tropism for neurons and large spatial spread after direct intracranial delivery^30^. This vector encoded the DREADD receptor hM4Di, fused to the fluorescent reporter mCherry to facilitate histological visualization. This gene was encoded downstream of a CaMKIIa promoter, which was used to target ATAC specifically to excitatory neurons^31^.

**Figure 2.**
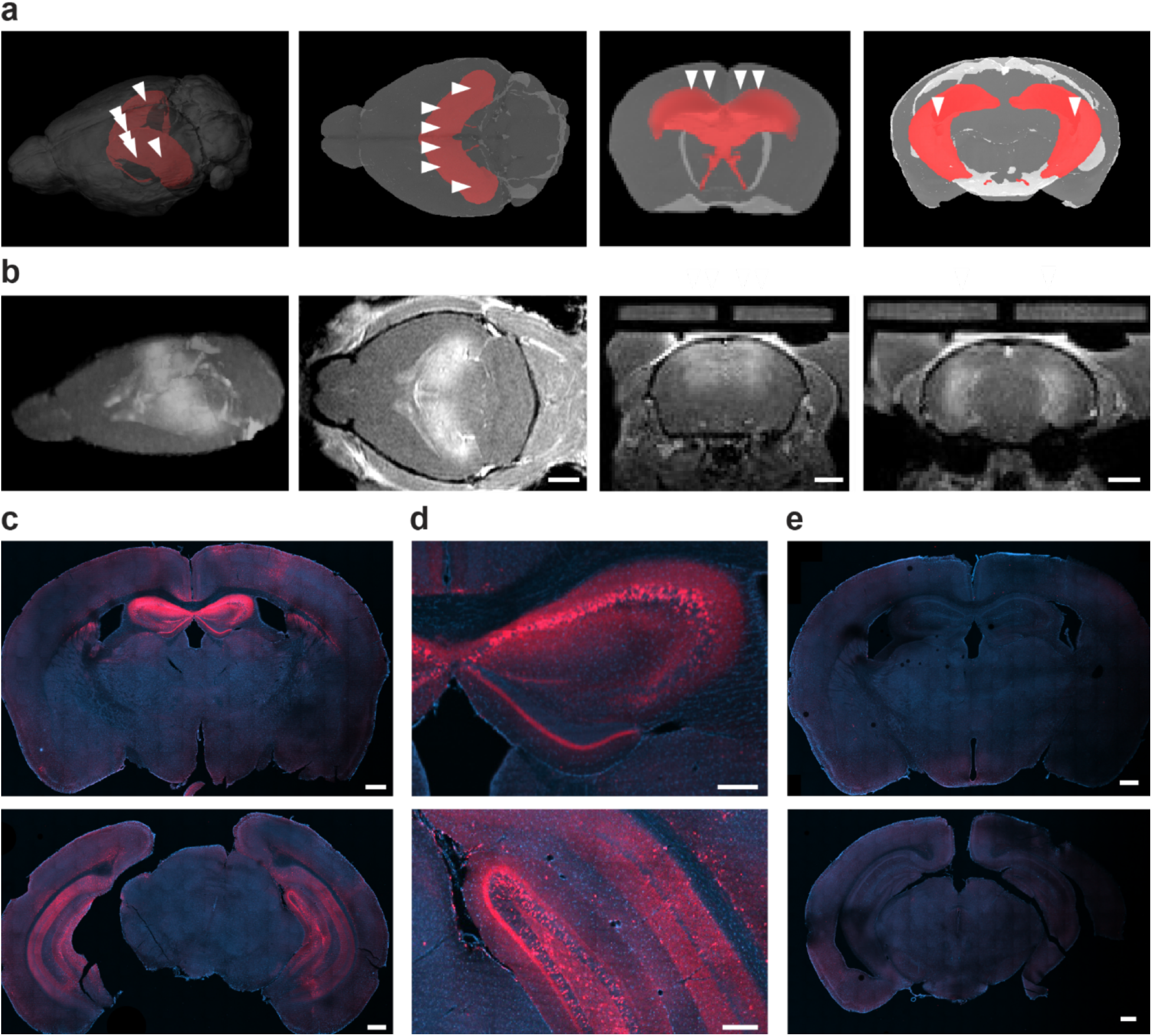
Blood-brain barrier opening and targeted expression of DREADD in the hippocampus. (**a**) Rendering of mouse brain with hippocampus highlighted in red and targeted locations of FUS-BBBO beams indicated with arrowheads. (**b**) Images from a representative T_1_-weighted MRI scan acquired immediately after FUS-BBBO, with brighter areas indicating relaxation enhancement from Prohance extravasation. Arrowheads indicate the locations of the FUS-BBBO targets. Scale bars 2 mm. (**c**) Representative brain section immunostained for hM4Di-mCherry (red) 6 weeks after FUS-BBBO and injection of AAV9 encoding this hM4Di-mCherry under the CaMKIIa promoter. The DAPI stain demarcates cell nuclei (blue). Scale bar, 500 μm. (**d**) Magnified view of the dorsal (top) and ventral (bottom) hippocampus showing widespread expression in molecular layers of the dentate gyrus, stratum orens, subiculum and granular cell layers of hippocampus. Scale bar, 200 μm. (**e**) Representative immunostaining result for hM4Di-mCherry in a mouse that received the same viral construct, but did not undergo FUS-BBBO. Scale bar, 500 μm. Histological images are representative of N=4 animals for each condition, and MRI images for N=24 animals.

After allowing 6–8 weeks for transgene expression, the targeted locations showed widespread hM4Di expression in histological sections, covering most hippocampal regions and a small segment of cortex above the dorsal hippocampus (Fig. 2, c). Expression was especially strong in the molecular and polymorph layers of dentate gyrus (DG), the stratum oriens and stratum radiatum of the CA1, CA2 and CA3 fields, as well as the pyramidal cells of the DG, CA2, and CA3 (Fig. 2, d). Expression was present broadly throughout the hippocampus (**Supplementary Fig. 1**). By comparison, mice that received an intravenous injection of the same AAV9 vector without FUS-BBBO showed essentially no fluorescent signal in these brain regions (Fig. 2, e), confirming that BBBO was required for gene delivery at the viral dose used in this study.

A quantitative comparison of expression in FUS-targeted areas across 5 mice was performed using mCherry fluorescence in cell bodies of granular cell layers, which allowed for a direct comparison of transfection efficiency between hippocampal regions. Our analysis showed that more than 50% of the cells in dorsal and ventral CA3 and dorsal CA2 were successfully transfected, and that ventral and dorsal DG contained 42% and 36% positive cell bodies, respectively (Fig. 3 a-b). Cortex and CA1 typically had lower transfection efficiencies, suggesting that these regions are less susceptible to transfection after BBBO than other hippocampal fields. As a representative non-targeted region, we looked for expression in the thalamus, which was shown in previous studies to be particularly susceptible to transfection following systemic delivery of AAV9^32^, and found no significant expression (<5%, Fig. 3 a-b). Full results of the statistical tests can be found in **Supplementary Table 1**.

**Figure 3.**
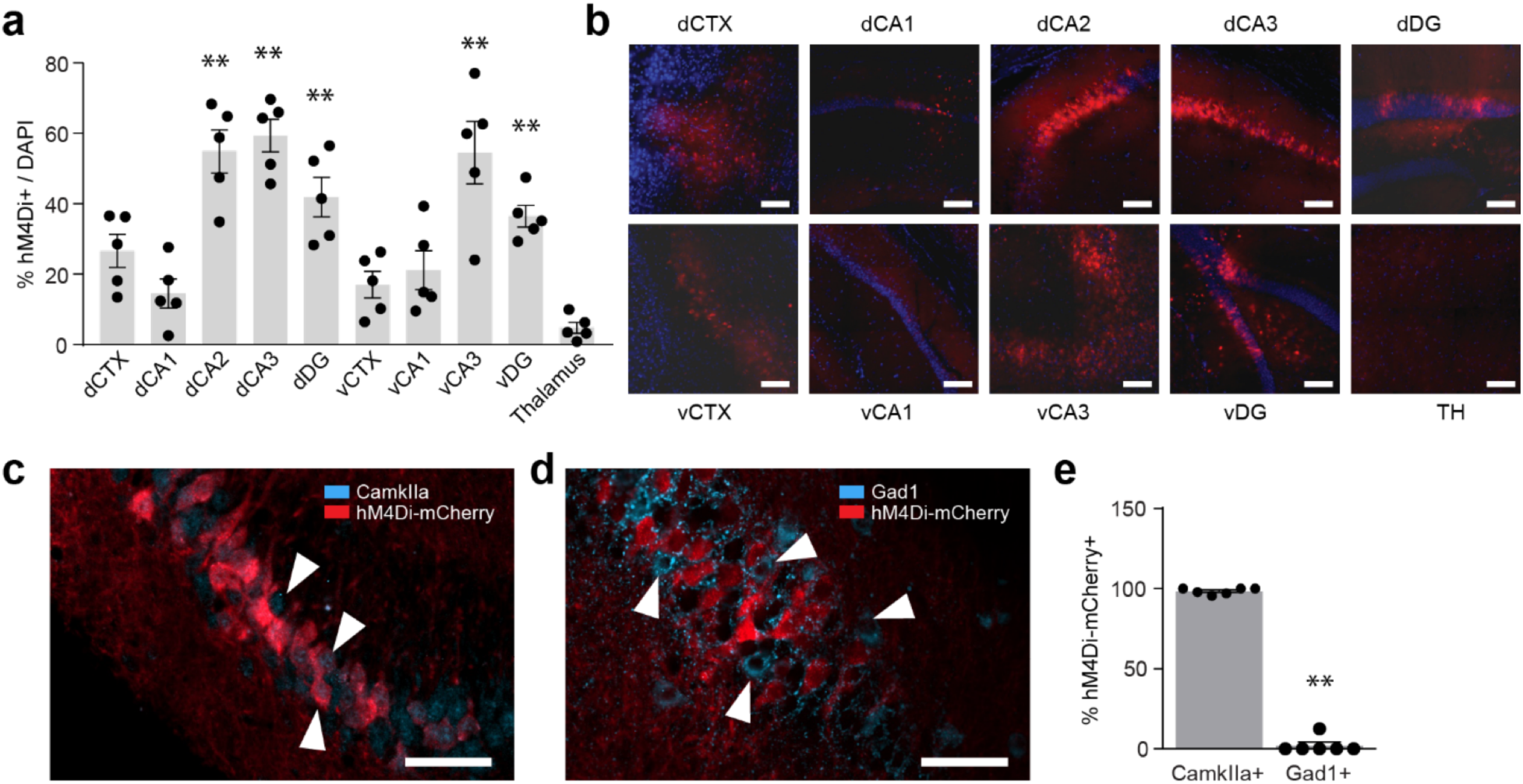
Spatial and cell type specificity of DREADD expression. (**a**) Percentage of cells bodies with detectable mCherry fluorescence in pyramidal layers of the hippocampus and overlaying FUS-targeted cortex, and thalamus as an untargeted negative control. The letters v and d indicate ventral and dorsal sites, respectively. n=5 mice; one-way ANOVA test compared to thalamus, F_(9, 40)_ = 13.89; p-value 8.7E-10, with Tukey-HSD post-hoc test; ** represents p<0.01. (**b**) Representative images of mCherry fluorescence (red) in each field. The DAPI stain (blue) marks cell nuclei. Scale bars represent 100 μm. (**c**) Representative co-immunostaining for hM4Di-mCherry and CaMKIIa. Arrowheads indicate cells positive for CaMKIIa. (**d**) Representative co-immunostaining for hM4Di-mCherry and Gad1. Arrowheads indicate cells positive for Gad1. (**e**) Percentage of DREADD-expressing cells in the CA2 region that are positively stained for CaMKIIa or Gad1, representing excitatory and inhibitory cells, respectively. n=6 mice; p<5E-9, two-tailed t-test assuming unequal variance. Scale bars in c-d are 50 μm. Bar graphs represent the mean ± SEM.

To evaluate the cellular specificity of genetic targeting, we compared expression of DREADDs in cells staining positively for CaMKIIa or Gad1, which serve as labels of excitatory and inhibitory neurons, respectively (Fig. 3, c-d). We found that 98.4 ± 0.8% of the cells expressing the DREADD receptor also stained positively for CaMKIIa, while only 2.08 ± 2.08% of these cells co-stained for Gad1, confirming selective targeting of our constructs to excitatory neurons (n=6, p<5E-9, heteroscedastic, two-tailed, t-test Fig. 3 e).

### Targeted stimulation of the hippocampus

Depending on the DREADD encoded in the AAV vector, ATAC can be used to either stimulate or inhibit targeted neurons. To first assess the ability of this technique to provide cell-specific activation, we targeted AAV9 carrying the excitatory DREADD hM3Dq-mCherry, under the CaMKIIa promoter, to the dorsal hippocampus using 4 FUS-BBBO sites. After allowing time for expression, we administered CNO intraperitonealy (IP), and 2.5 hours later perfused the mice to histologically monitor the expression of the activity-dependent gene product c-Fos in the dorsal CA3 region of the hippocampus (Fig. 4, a). We found that cells positive for hM3Dq expression were 5.8 times more likely to exhibit c-Fos staining than cells not expressing hM3Dq (Fig. 4, b, c, n=6, p<0.001, paired t-test), indicating that systemic chemogenetic treatment allows ATAC-targeted neurons to be selectively activated several weeks after the spatial targeting procedure.

**Figure 4.**
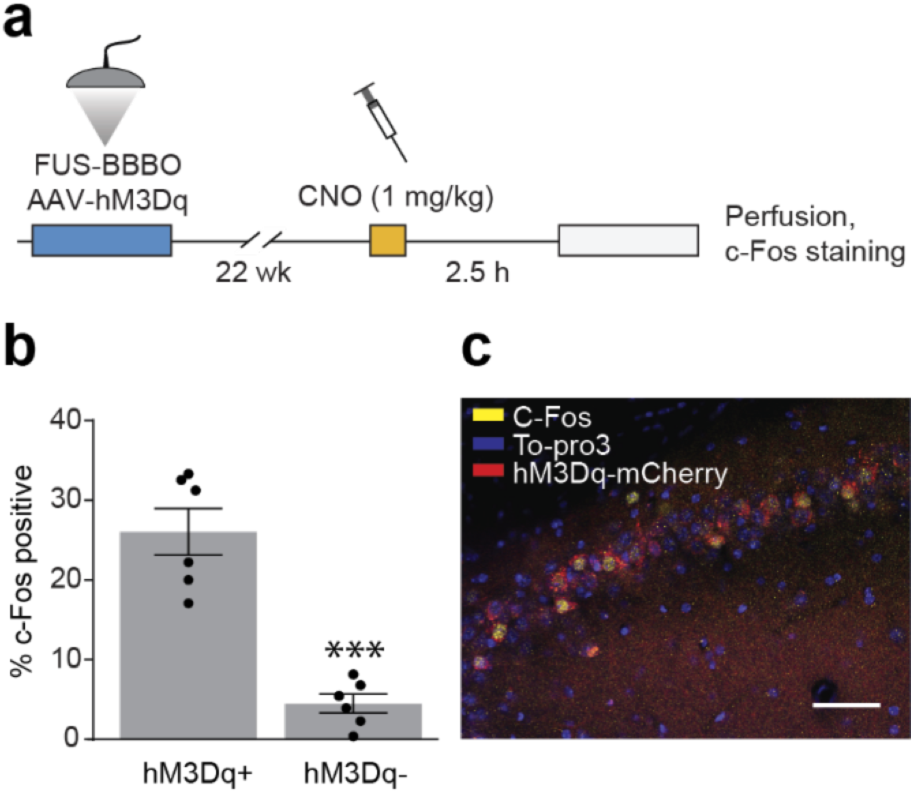
ATAC with excitatory DREADDs results in neuronal activation. (**a**) Excitatory DREADD activation protocol. FUS-BBBO and an IV injection of AAVs was followed by a period of expression, and an IP injection of CNO in saline (1 mg/kg). 2.5 hours later, mice were perfused and their brains were extracted for histological evaluation. (**b**) Fraction of granular cells in the CA3 field of the hippocampus staining positively for c-Fos after CNO administration, as a function of whether the cells are positive or negative for hM3Dq-mCherry. (***, p<0.001, two tailed t-test, assuming unequal variance, n=6 independently targeted hemispheres in n=3 mice). (**c**) Representative immunohistology image of CA3 with c-Fos (yellow), hM3Dq-mCherry (red) and nucleus (To-pro3, blue) staining. Scale bar, 50 μm. Bar graphs represent the mean ± SEM.

### Targeted inhibition of memory formation

To assess the ability of ATAC to inhibit targeted neurons, and to test the functionality of this technology in a behavioral paradigm, we used FUS-BBBO to target inhibitory DREADDs (hM4Di) to ventral and dorsal hippocampus (Fig. 5, a), and assessed the impact of CNO administration on the formation of contextual fear memories. This well-established behavioral paradigm has been used in studies of anxiety, phobias, PTSD and fear circuitry^33, 34^. Fear conditioning has also served as a testing ground for other novel neuromodulatory techniques^35,36^. Since the hippocampus plays an essential role in memory formation, we hypothesized that inhibiting it noninvasively using the ATAC strategy would reduce the ability of mice to form fear memories.

**Figure 5.**
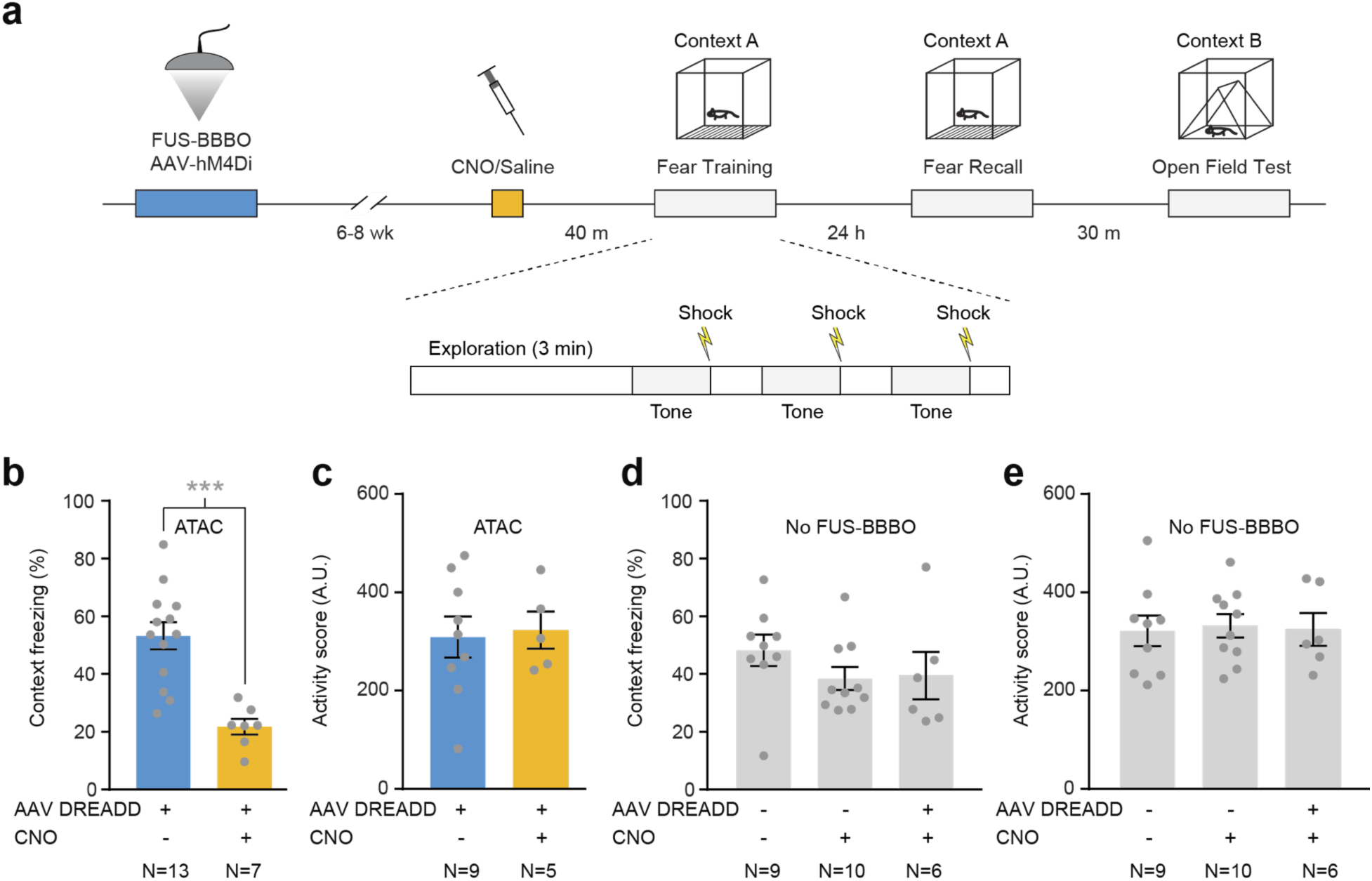
Inhibition of fear memory formation using ATAC. (**a**) Illustration of the ATAC and fear conditioning protocol. 6–8 weeks after FUS-BBBO and administration of AAV-DREADD (hM4Di-mCherry), mice were injected with CNO or saline, then placed in a fear conditioning chamber with an electrified floor. After 3 minutes of free exploration, the mice received 3 × 30-second tones (80 dB) paired with an electric shock during the last 2 s of the tone (0.7 mA) There was a 1-minute interval between tones. 24 h after training, the mice were placed in the same chamber and allowed to explore for (8 min 40 s). 30 min later, the mice were placed in a different context for a 3-minute open field test. (**b**) Percentage of time spent freezing in the fear recall context (Context A) for ATAC mice treated with CNO or saline (p < 2E-5, two-tailed t-test assuming unequal variance). (**c**) Exploratory activity score in the non-fear context (Context B) for ATAC mice treated with CNO or saline (p=0.81, two-tailed t-test assuming unequal variance). (**d**) Percentage of time spent freezing in the fear recall context (Context A) for wild-type mice treated with CNO or saline, or CNO-treated mice that received IV injection of AAV-DREADD without FUS-BBBO (no effect found via one-way ANOVA, F_(2,22)_=1.005, p=0.38). (**e**) Exploratory activity score in the non-fear context (Context B) for wild-type mice treated with CNO or saline, or CNO-treated mice that received IV injection of AAV-DREADD without FUS-BBBO (no effect found via one-way ANOVA, F_(2,22)_=0.04, p=0.96). Bar graphs represent the mean ± SEM. N provided under each bar indicates number of mice.

Coverage of the hippocampus was achieved with FUS-BBBO applied to 6 sites (Fig. 2), accompanied by intravenous administration of AAV9 containing hM4Di-mCherry under the CaMKIIa promoter. Two groups of control mice were either completely untreated or received intravenous virus without FUS-BBBO. After 6–8 weeks, the mice underwent a fear conditioning protocol (Fig. 5a). In this protocol, the mice are placed in a unique environment (defined by chamber shape, lighting, smell and sound; Context A in Fig. 5a) while receiving three electric foot shocks, causing them to associate this environment with the noxious stimulus in a process known as context fear conditioning. 40–60 minutes before undergoing this protocol, the mice received injections of either CNO or saline to test the ability of targeted inhibition of ATAC-treated hippocampal neurons to reduce fear formation.

24 hours after conditioning, contextual fear recall was tested by placing mice in the same context and measuring freezing, an indication of fear^37^ (Fig. 5a). ATAC mice treated with saline during conditioning froze 53.2% of the time, indicating robust fear recall. By contrast, ATAC mice that received CNO before conditioning froze only 21.8 % of the time – a more than 2-fold reduction in fear memory (p<2E-5; n=13 and 7, heteroscedastic, two-tailed, t-test). (Fig. 5b). Comparing these two FUS-treated conditions to each other allowed us to evaluate the efficacy of ATAC while accounting for any potential behavioral effects caused by the FUS treatment itself^38,39^. To confirm that the effect of activation of DREADDs was specific to fear formation, and did not affect basic exploratory behavior, we monitored treated and untreated mice in an open field test. One day after fear conditioning, the mice were placed in a novel environment (Context B in Fig. 5a), which they were allowed to explore freely for 3 minutes. Their total mobility, quantified automatically from video recordings, was unchanged between CNO and saline groups (Fig. 5c).

Additional controls showed that activation of any DREADDs potentially expressed in peripheral organs (in the absence of FUS-BBBO), or CNO treatment alone (in wild-type mice) did not result a reduction in context fear relative to untreated controls (Fig. 5d). Exploratory behavior was likewise unaffected (Fig. 5e).

To confirm that the effect of ATAC treatment was specific to inhibiting memory formation as opposed to sensation of stimuli such as pain, we paired each foot shock with an audible tone to produce an association between the tone and the shock in a process known as cued conditioning (**Supplementary Fig. 2a**). This process takes place immediately during training, and is not expected to be affected by inhibition of memory-forming regions of the hippocampus^36^. As expected, cued freezing measured at the end of the training session was unaffected by CNO treatment (**Supplementary Fig. 2, b-c**), indicating that ATAC-treated mice were not compromised in their ability to experience salient sensory stimuli.

### Intersectional ATAC in transgenic animals

In addition to its potential therapeutic applications, ATAC may facilitate the study of neurological and psychiatric disease mechanisms in animal models by making it possible to modulate disease-related spatially defined neural circuits without surgery. A complementary resource for such studies is the large number of transgenic mouse and rat lines available with cell type-specific expression of the CRE recombinase. The delivery of a viral vector encoding any gene of interest in a CRE-dependent configuration allows the expression of this gene to be confined to the CRE-expressing cells in that animal^25^. To test whether ATAC could be used in combination with a CRE mouse line to provide noninvasive spatial and cell-type targeting of neuromodulation, we used FUS-BBBO to deliver a CRE-dependent DREADD construct into TH-CRE transgenic mice^40^. These animals express the CRE recombinase in tyrosine hydroxylase-positive dopaminergic neurons in the midbrain, especially in the substantia nigra pars compacta (SNc) and the ventral tegmental area (VTA). These regions are researched extensively in models of Parkinson’s disease^41^, addiction and reward^42^ and have previously been used to validate new neuromodulation techniques^43^. Due to their locations deep within the brain and their small size, surgical access to these sites is difficult, and a noninvasive approach could reduce surgical damage seen along the needle tract while providing spatial selectivity.

To establish the feasibility of intersectional ATAC in a CRE mouse line, we used FUS-BBBO to spatially target CRE-dependent^44^ DIO-Syn1-hM3Dq-mCherry encoded in AAV9 to the midbrain on a single side of the brain (Fig. 6, a), then tested our ability to activate TH-positive neurons in this region with CNO by imaging c-Fos accumulation (Fig. 6, b). FUS-BBBO applied to the midbrain resulted in BBB opening that partially overlapped with the expected location of the SNc/VTA (Fig. 6, c). Subsequent immunofluorescent imaging of brain sections revealed mM3Dq expression specific to the SNc/VTA region at the FUS-BBBO site (Fig. 6, d-e). We then tested the functionality of our DREADD receptor by staining for c-Fos positive nuclei at the site of FUS-BBBO and the contralateral region. Among TH-positive neurons, we found a 7.3-fold increase in activation on the side targeted by the ATAC treatment (Fig. 6, f; n=5 mice, p<1.1E-3, paired t-test), demonstrating spatially selective neuromodulation in a CRE model.

**Figure 6.**
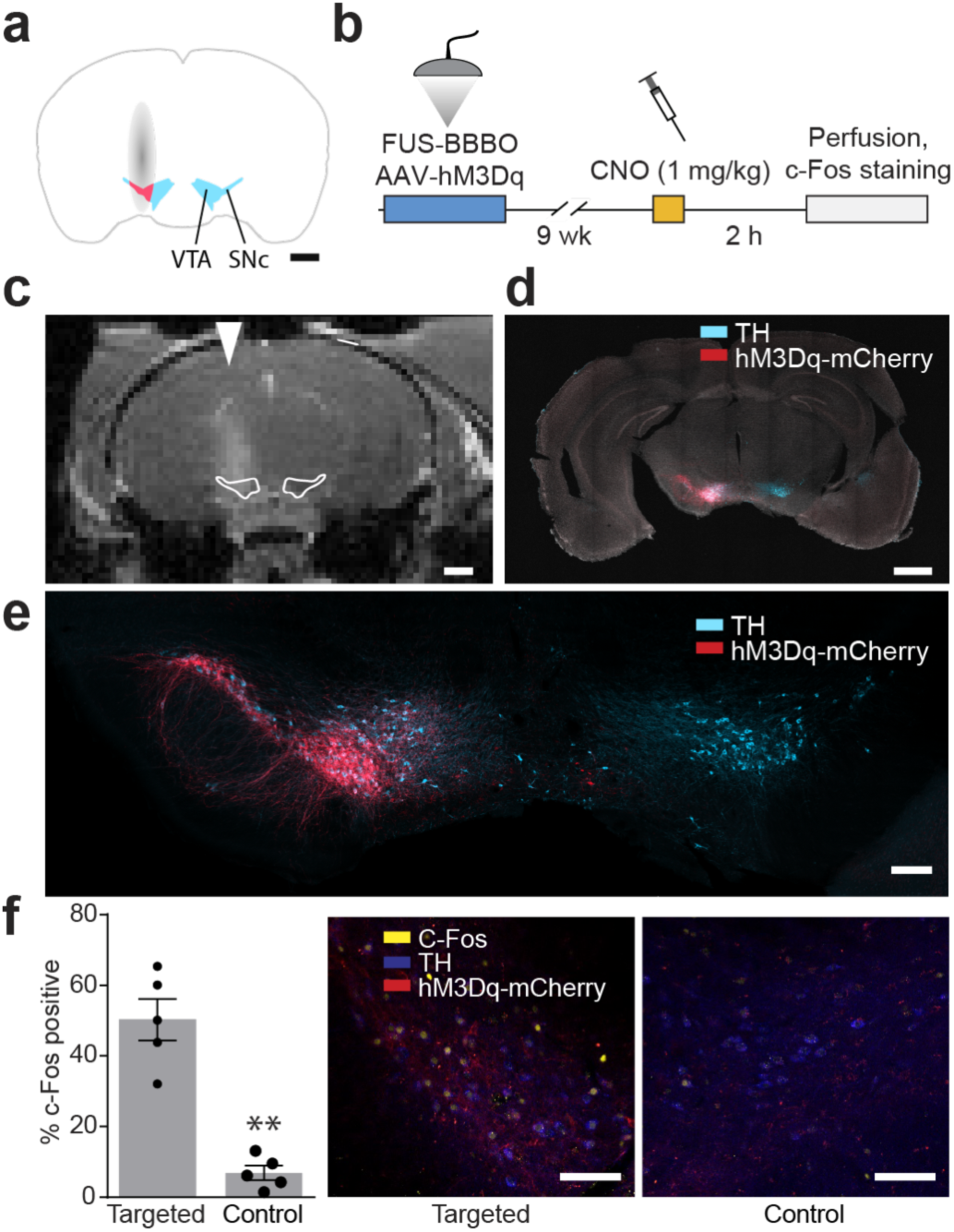
Intersectional ATAC in the midbrain of CRE transgenic mice. (**a**) Illustration of intersectional attack experiment. FUS-BBBO (grey) was used to target AAV encoding DIO-Syn1-hM3Dq-mCherry unilaterally to the midbrain of TH-CRE mice. Approximate locations of the TH-positive SNc/VTA and FUS target region shown in cyan and pink, respectively. Scale bar is 1 mm. (**b**) Protocol for c-Fos induction. After a period of expression, mice received an IP injection of CNO (1 mg/kg), and 2 hours later were perfused and their brains extracted for histological evaluation. (**c**) Representative T_1_-weighted MRI scan indicating the site of BBBO (representative of 7 mice). Outlines show the approximate location of SNc/VTA. The arrowhead indicates the lateral targeting of FUS. Scale bar is 1 mm. (**d**) Immunostaining for hM3Dq-mCherry (red) and TH (blue), counterstained with DAPI (white). 5 mice were evaluated with similar results. Scale bar is 1 mm. (**e**) Magnified view of VTA/SNc area in (d). Scale bar is 200 μm. (**f**) Quantification of activated (c-Fos-positive), TH-positive neurons in the ATAC-targeted SNc/VTA region after treatment with CNO, compared to contralateral control (p < 1.1E-3, paired, two-tailed, t-test, n=5), together with representative histology images of the targeted and contralateral brain regions stained for c-Fos (yellow), TH (blue) and hM3Dq-mCherry (red). Scale bar is 100 μm. Bar graphs represent the mean ± SEM.

### Tolerability of ATAC by brain tissue

Finally, to confirm that ATAC treatment is well tolerated by brain tissue, we examined hematoxylin-stained brain sections from 14 mice with a total of 84 FUS-BBBO sites. Consistent with previous findings^45^, the majority of these sites (71.4 %) had normal histology (**Supplementary Fig. 3, a-b**). In the remaining FUS-targeted sites we found small histological features with mean dimensions of 115 μm by 265 μm, which were not visible on sections ± 300 μm away from the site (**Supplementary Fig. 3, b-c**). The average calculated volume of these features was 0.0027 ± 0.0007 mm^3^. This represents less than 0.1% of the typical FUS-BBBO site, which has a volume of 2.81 ± 0.51 mm^3^ (average of n=7 sites quantified by MRI) and 0.01% of the mouse hippocampus (volume, 26 mm^3^)^46^. By comparison, the brain volume displaced by a typical needle during invasive viral injections is approximately 0.23 mm^3^ (**Supplementary Fig. 3, d**). These results are consistent with the normal performance of ATAC-treated mice in behavioural tests and the ability of ATAC-treated regions to become chemogenetically activated and express c-Fos. In future translational studies, this safety profile could be further improved with feedback-controlled FUS-BBBO^47^.

## DISCUSSION

Taken together, our results establish ATAC as a paradigm for noninvasive neuromodulation with a unique combination of spatial, cell-type and temporal specificity. This paradigm holds several advantages over existing techniques for both research and potential clinical applications. Compared to intracranial injections for viral gene delivery, which are invasive and often require multiple brain penetrations to cover the desired area, FUS-BBBO enables comprehensive transduction of an entire brain region in a single session with minimal tissue disruption, and can easily be scaled to larger animals and humans. While recent developments in AAV vectors also enable some variants to cross the BBB on their own^32^, they do so without spatial selectivity.

Compared to emerging ultrasonic neuromodulation techniques in which ultrasound directly activates or inhibits brain regions or releases local neuromodulatory compounds^48–55^, ATAC does not require an ultrasound transducer to be mounted on the subject during modulation. After transduction and expression of chemogenetic receptors in a genetically defined subset of cells at the FUS-targeted site, neuromodulation is conveniently controlled using an orally bioavailable drug. The fact that a single FUS-BBBO session is required should also minimize the potential for non-specific cellular-level effects seen after multiple FUS-BBBO treatments^56,57^.

In our behavioral proof of concept, a single injection of CNO several weeks after the FUS-BBBO procedure resulted in a 2.4-fold reduction in fear memory formation without any effects on normal exploratory behavior. In addition, both the cell types modulated in the chosen brain region and the polarity of the modulation can be chosen precisely using cell type-specific promoters and excitatory or inhibitory receptors. Finally, we showed that ATAC is compatible with intersectional genetic targeting in transgenic animals, making it potentially useful in a wide variety of basic and disease model studies.

The ATAC paradigm could be made more powerful with improvements in each of its components: FUS-BBBO, AAV vectors, cell-specific promoters, chemogenetic receptors and ligands. For example, the BBBO procedure can be improved using real-time monitoring of bubble cavitation to maximize molecular delivery while minimizing the possibility of damage^47^. In scale-up to larger animals and humans, FUS also benefits from the use of phased array sources and image-based aberration correction^9,58–61^. Additional work is needed to make AAV vectors more efficient to reduce their required dose, and to make compact cell-type specific promoters that work robustly in primates^24^. Finally, ongoing studies of the pharmacokinetics of CNO and clozapine^62^ will enable the optimization of ligand dosing for DREADD activation or motivate the use of available alternative ligands^63^ or chemogenetic receptors^64,65^. These improvements will facilitate the development and translation of ATAC as a paradigm for precise noninvasive control of neural circuits.

## MATERIALS AND METHODS

### Animals

C57BL6J mice were obtained from JAX laboratories. Transgenic TH-CRE mice were obtained from a Caltech’s internal colony, and were originally generated^40^ at Uppsala University, Sweden. Animals were housed in 12 hr light/dark cycle and were provided water and food ad libitum. All experiments were conducted under a protocol approved by the Institutional Animal Care and Use Committee of the California Institute of Technology.

### FUS-BBBO and Viral Delivery

Male, 13–18 week old C57BL6J mice were anesthetized with 2% isoflurane, the hair on their head removed with Nair depilation cream and then cannulated in the tail vein using a 30 gauge needle connected to PE10 tubing. The cannula was then flushed with 10 U/ml heparin in sterile saline (0.9% NaCl) and affixed to the mouse tail using tissue glue. Subsequently, mice were placed in the custom-made plastic head mount and imaged in a 7T MRI (Bruker Biospec). A FLASH sequence (TE=3.9 ms, TR=15 ms, flip angle 20 degrees) was used to record the position of the ultrasound transducer in relation to the mouse brain. Subsequently, mice were injected via tail vein with AAV9 encoding DREADDs (pAAV-CaMKIIa-hM4D(Gi)-mCherry or pAAV-CaMKIIa-hM3D(Gq)-mCherry (gifts from Bryan Roth) and immediately after with Definity microbubbles (Lantheus) and Prohance (Bracco Imaging) dissolved in sterile saline. Within 30 seconds, mice were insonated using an 8-channel focused ultrasound system (Image Guided Therapy, Bordeux, France) driving an 8-element annular array transducer with diameter of 25mm, coupled to the head via aquasonic gel. The ultrasound parameters used were 1.5 Mhz, 1% duty cycle, and 1 Hz pulse repetition frequency for 120 pulses. For each FUS site, Definity and Prohance were re-injected before insonation. The total dose of Prohance injected was 0.5 μmoles/g. After FUS-BBBO, the mice were imaged again using the same FLASH sequence to confirm opening of the BBB and appropriate targeting. Immediately afterwards, mice were placed in the home cage for recovery. The TH-CRE animals, aged 18 weeks, were subjected to FUS-BBBO using the same protocol, and using the same dose of AAV9, as for C57BL6J mice. All TH-CRE animals were females.

MRI images were analyzed using imageJ measurement function. To estimate size of the BBB opening, we used a single FUS-beam using standard parameters. The hyperintense area from Prohance extravasation was delineated manually and the dimensions of minor and major axis recorded for n=7 animals. The volumes were calculated assuming ellipsoid shape. For MRI intensity calculation, the 4 sites of FUS-BBBO in dorsal hippocampus were delineated manually and average signal intensity calculated within the region for each mouse. The result was then divided by a mean signal intensity in an untargeted thalamus 1.5–2mm below hippocampus.

### C-Fos activation and immunostaining

C57BL6J male mice of 13 weeks of age underwent FUS-BBBO to administer AAV9 carrying hM3Dq-mCherry into the hippocampus. Subsequently, mice were housed singly to reduce background C-fos expression. After 22 weeks of expression, mice received an IP injection of 1 mg/kg CNO in sterile saline, and were returned to home cages. After 150 minutes, mice were anesthetized using Ketamine/Xylazine solution (80 mg/kg and 10 mg/kg, respectively, in PBS) and perfused with a cold PBS/Heparin (10 U/ml), and immediately afterwards with 10 % Neutral Buffered Formalin (NBF). Their brains were extracted and post-fixed in 10% NBF for at least 24 hours. Brain sections (50 μm) were obtained using a VF-300 compresstome (Precisionary Instruments). Subsequently, sections were blocked in 10% Normal Donkey Serum and 0.2% Triton-X solution in PBS for 1 hr in room temperature, and immunostaining with primary antibody was performed using a goat anti-c-Fos antibody (SC-253-G, SCBT, Santa Cruz, CA) in 10% Normal Donkey serum and 0.2% Triton-X, overnight at 4 °C. Afterwards, sections were washed three times in PBS and incubated with a secondary donkey anti-goat antibody conjugated to Alexa-488 (A-11055, Thermofisher). For activation of hippocampus, the histological evaluation was performed by an observer blinded to the identity (hM3Dq positive, or negative) of granular layer nuclei in dCA3. The expression status of the neurons was determined after the scoring of c-Fos positivity. The activation of TH neurons was evaluated by an observer blinded to the presence of FUS-BBBO targeting at a given site.TH-CRE mice expressed hM3Dq for 9 weeks after FUS-BBBO, and then were given 1 mg/kg CNO. After 2 h, they were anesthetized with ketamine/xylazine (80/10 mg/kg in PBS) and perfused using cold heparine/PBS (10 u) and then 10% NBF. One region of interest in 1 out of 6 TH-CRE mice was damaged during sectioning and the mouse couldn’t be included in c-Fos evaluation.

### Gene expression evaluation

To visualize DREADD expression across brain regions, we used immunostaining with a polyclonal rabbit anti-mCherry antibody (PA534974, Thermofisher), a polyclonal goat anti-CaMKIIa antibody (PA519128, Thermofisher) and a polyclonal goat anti-Gad67 (Lifespan, 103220–296) antibody in 10% Normal Donkey Serum (NDS, D9663–10ML, Sigma-Aldrich) and 0.2% Triton-X in PBS, overnight at 4 °C. The TH expression was evaluated using an anti-TH chicken antibody (TYH, Aves lab) incubated in normal Goat Serum (NS02L-1ML, Sigma-Aldrich) and 0.2% triton-x in PBS at 4 °C, overnight. Secondary antibodies were donkey anti-rabbit conjugated to Dylight-650 (#84546, ThermoFisher), donkey anti-goat conjugated to Dylight-488 (SA510086) and goat anti-chicken conjugated to Alexa 488 (A-11039, ThermoFisher). Secondary antibodies were incubated in 10% NDS/0.2% triton-x in PBS for 4 h at room temperature. For quantitative comparison of expression levels between various regions of the hippocampus, we used mCherry fluorescence localized to cytoplasmic compartments and counted the number of cells in the pyramidal layers of hippocampus showing detectable fluorescence. Cells were co-stained with a nuclear stain (DAPI) to allow delineation of nuclei and surrounding cytoplasmic regions. Cells that showed mCherry fluorescence surrounding the nucleus for at least 50% of its circumference were counted as positive. All the images were background-normalized to allow for comparable evaluation of expression. The inter-experimenter variability was determined for two different researchers (B.L., J.O.S), for n=6 samples, with the difference in means smaller than 2.5% (mean = 42.5% vs 41.5%, p=0.92, heteroscedastic, two-tailed, t-test).

### Behavioral testing

Behavioral studies for fear conditioning were performed in sound-attenuated fear conditioning chambers (30 × 25 × 25 cm, Med Associates). Animals were trained and tested for context fear in Context A, which comprised a staggered wire grid floor, white light, 5% acetic acid for scent and no background noise. Locomotor testing was performed in Context B, which was differentiated from Context A by chamber shape, floor, illumination, odor, background noise and room location. Animal activity was recorded and quantified using Video Freeze software (Med Associates). For cued training, the tone was 80 dB and 30s. **Fear conditioning:** Mice were injected with CNO (10 mg/kg, IP) or saline (IP), and after 40–60 minutes underwent context and cued fear conditioning in Context A. A 3-minute initial baseline period was followed by 3 × 30-s presentations of a tone co-terminated with a 2-s foot shock (0.7 mA), with inter-trial interval of 60 s. After the trials, the mice remained in the context for an additional 60 s, after which they were transported back into the vivarium. After 24 h, mice were placed in Context A to record context fear for the duration of training (8 min. and 40 s). **Exploratory behavior analysis:** Between 30 and 45 minutes after the context fear test, mice were transported to another room, placed in Context B and allowed to explore the chamber for 3 minutes while their activity was recorded. Due to automated data acquisition and evaluation, no blinding was necessary. **Fear conditioning analysis**. Mice were recorded using automated near-infrared video tracking in the fear conditioning chamber using VideoFreeze software. Mouse motion was measured using the activity score, from a video recording at 30 frames/s, with the motion threshold set at 18 activity units (standard value set in software). Freezing was defined as an activity score below 18 units for at least 1 s. Average freezing in the context test was scored over the whole trial. Due to automated data acquisition and evaluation, no blinding was necessary. **Exclusions:** mice were excluded from statistical analysis if their histologically determined DREADD expression was below 30% of cell bodies in dorsal CA3 region of the hippocampus. This threshold was chosen based on previous studies showing that behavioral effects generally require modulation of at least 30% of the neurons in a targeted region^66,67^ and dorsal CA3 being the most robustly transfected hippocampus region. The resulting analyzed groups had identical levels of expression (55.1% for Saline and 60.5% for CNO groups, p=0.26, heteroscedastic,two-tailed, t-test). In analyses including all mice, we found that DREADD expression in dorsal CA3 correlated with the formation of context fear memories in mice treated with CNO (r=0.62, n=11) but not in mice receiving saline (r=0.14, n=14) (**Supplementary Fig. 4, a**). Even without excluding the four mice who had expression below 30%, a direct comparison between ATAC mice treated with CNO and saline showed a statistically significant reduction of context fear (53.2 vs 34%, n=13, 11; p<0.02; heteroscedastic, two-tailed, t-test). Variability in gene expression may have been due to poor intravenous injections of virus during the FUS-BBBO procedure, since we found no difference between these mice in T_1_ MRI signal enhancement post FUS-BBBO (**Supplementary Fig. 4b**).

### Statistical analysis

Data was analyzed using either two-tailed t-test with unequal variance (when two samples were compared and when data was deemed normal with Shapiro-Wilk test) or one-way ANOVA with a Tukey HSD post-hoc test (when more than two samples were compared).All data with p<0.05 were considered significant. Error bars used throughout the study represent standard error of mean (SEM). “*” Corresponds to p<0.05, “**” to p<0.01 and “***” to p<0.001. All data was tested for normality using Shapiro-Wilk test. Samples with two conditions and non-normal distributions were tested by a nonparametric test (Mann-Whitney). All central tendencies reported are averages.

### Histological analysis

Analysis of the FUS-BBBO safety was performed using hematoxylin staining and autofluorescence. All of the vibratome sections (50 micron) within the hippocampus were imaged under 10x objective under microscope to identify potential lesions (n=14 mice). The sections showing largest anomalies were then observed in a greater magnification (20x objective). The FUS-induced lesions were autofluorescent and fluorescence microscopy was used for measurements. The volumes were calculated assuming ellipsoid shape of the damage, with maximum diameters within a section used for major and minor axes. The volume of lesions was calculated using ellipsoid volume formula (v = 4/3 × π × (width/2)^2^ × length/2). To confirm the anatomy of the lesions, hematoxylin staining was performed: vibratome sections were stained for 30–45 seconds in 20% Gill no. 3 hematoxyllin, followed by a brief wash in PBS and 5 second dip in RapidChrome blueing solution (Thermofisher). Each section was then washed twice in PBS and mounted in a water-based medium (ProLong Gold, Thermofisher).

### Illustrations

The structure of AAV9^68^ in Figure 1 has been rendered using QuteMol^69^. The 3D rendering of hippocampus in Figure 2 was generated using Rhinoceros 3D software with models obtained from 3D brain atlas reconstructor^70^ and waxholm space dataset^71^.

## ACKNOWLEDGEMENTS

The authors thank Moriel Zelikowsky for insightful discussions and assistance with the design of fear conditioning experiments. We thank Erik Dumont, Russ Jacobs, Arnab Mukherjee and George Lu for helpful and constructive discussions and Reed McCardell for assistance with initial experiments. We thank Viviana Gradinaru and Ben Deverman for helpful discussions and providing TH-CRE mice. We appreciate the help and assistance of the UCLA’s Translational Pathology Core Laboratory with imaging of some of the microscopy samples and Caltech’s Office of Laboratory Animal Research for help with rodent husbandry. This research was supported by the Heritage Medical Research Institute (M.G.S.), DARPA (M.G.S., J.S.) and the Jacobs Institute for Molecular Medicine (M.G.S., J.S.).

## AUTHOR CONTRIBUTIONS

J.S. and M.G.S. conceived and planned the research. J.S. performed the *in vivo* experiments. J.S. B.L. A.L.G. and D.M. performed histological experiments. J.S. and B.L. analyzed data. J.S., and M.G.S. wrote the manuscript with input from all other authors. M.G.S. supervised the research.

